# Quantitative Models of Phage-Antibiotics Combination Therapy

**DOI:** 10.1101/633784

**Authors:** Rogelio A. Rodriguez-Gonzalez, Chung-Yin Leung, Benjamin K. Chan, Paul E. Turner, Joshua S. Weitz

## Abstract

The spread of multi-drug resistant (MDR) bacteria is a global public health crisis. Bacteriophage therapy (or “phage therapy”) constitutes a potential alternative approach to treat MDR infections. However, the effective use of phage therapy may be limited when phage-resistant bacterial mutants evolve and proliferate during treatment. Here, we develop a nonlinear population dynamics model of combination therapy that accounts for the system-level interactions between bacteria, phage and antibiotics for *in-vivo* application given an immune response against bacteria. We simulate the combination therapy model for two strains of *Pseudomonas aeruginosa*, one which is phage-sensitive (and antibiotic resistant) and one which is antibiotic-sensitive (and phage-resistant). We find that combination therapy outperforms either phage or antibiotic alone, and that therapeutic effectiveness is enhanced given interaction with innate immune responses. Notably, therapeutic success can be achieved even at sub-inhibitory concentrations of antibiotics, e.g., ciprofloxacin. These *in-silico* findings provide further support to the nascent application of combination therapy to treat MDR bacterial infections, while highlighting the role of innate immunity in shaping therapeutic outcomes.

## I. INTRODUCTION

Multi-drug resistant bacterial infections are a threat to global health. The World Health Organization (WHO) has reported that drug-resistant tuberculosis alone kills 250,000 people each year^1^. Moreover, the United States Centers for Disease Control and Prevention (CDC) have reported 23,000 deaths each year attributed to drug-resistant pathogens, while their European counterparts have reported 25,000 deaths each year resulting from drug-resistant infections^2,3^. The WHO has identified and prioritized twelve MDR pathogens^1^ in order to guide efforts toward the development of new antimicrobial treatments. The Gram-negative bacteria *Pseudomonas aeruginosa* (*P.a.*) has been identified as a critical priority by the WHO^1^.

Bacterial viruses (i.e., bacteriophage or “phage”) represent an alternative approach to treat MDR bacterial infections. Phage lysis of bacteria cells can drastically change bacterial population densities. In doing so, phage exert a strong selection pressure on the bacterial population. As a result, phage-resistant mutants can appear and become dominant^4–6^, whether via surface-based resistance^4,7^ or intracellular mechanisms^8^. The possibility that phage therapy may select for phage-resistant bacterial mutants has increased interest in identifying strategies to combine phage with other therapeutics, e.g., antibiotics^4,6,7,9–11^. However, the realized outcomes of combination strategies are varied, ranging from successes *in vitro*^9^ and *in vivo*^4,11^ to failure given *in vitro* settings^6^.

In many cases, the mechanism(s) underlying potential phage-antibiotic interactions are unknown. There are exceptions, for example, *Escherichia coli* phage TLS and U136B infect the bacterium by attaching to the outer membrane protein, TolC, which is part of the AcrAB-Tolc efflux system^12,13^. It has been shown that phage TLS selects for *tolC* mutants that are hypersensitive to novobiocin^13^. Moreover, TolC has been identified as a phage receptor in other Gram-negative pathogens^14,15^, giving further support to the combined use of phage and antibiotics. Similarly, the phage OMKO1 may be able to use multiple binding targets to infect *P.a.*, including the type-IV pilus and the multidrug efflux pump, MexAB/MexXY^7^, both mechanisms can result in selection against drug resistance.

The ability of phage OMKO1 to select against drug resistance in *P.a.* suggests that a combination treatment of *P.a.* with phage OMKO1 and antibiotics can lead to an evolutionary trade-off between phage and antibiotic resistance^7,11^. Phage-resistant mutants can show impairments of the multidrug efflux pump, MexAB/MexXY^7^, such as reduced functionality (or loss) of outer membrane porin M, OprM. This protein is part of the efflux pump complex and may act as a cell receptor of the phage OMKO1. Mutations in the gene encoding OprM can impair phage infection and restore the sensitivity to some classes of antibiotics, including ciprofloxacin^7^. Such an evolutionary trade-off may be leveraged clinically to limit the spread of resistance to phage and antibiotics.

Therapeutic application of phage and antibiotics *in vivo* necessarily involves interactions with a new class of antimicrobial agents: effector cells within the immune system. Recent work has shown that phage and innate immune cells, specifically neutrophils, combine synergistically to clear otherwise fatal respiratory infections in which neither phage nor the innate immune response could eliminate alone^5^. This ‘immunophage synergy’ is hypothesized to result from density-dependent feedback mechanisms^16^. Phage lysis decreases bacterial densities such that the activated immune response can clear bacteria; without phage the bacterial densities increase to sufficiently high levels that are outside the range of control by immune cells. However, the potential role of the innate immune response in the context of phage-antibiotic combination therapy remains largely unexplored.

Here, we develop and analyze a mathematical model of phage-antibiotic combination therapy that builds on the synergistic interactions between phage, antibiotic, and immune cells. In doing so we extend a mathematical model of immunophage synergy^16^ to take into account the pharmacodynamics and pharmacokinetics of an antibiotic, e.g., ciprofloxacin. At the core of the combination therapy model is its multiple-targeting approach, the phage target phage-sensitive (antibiotic-resistant) bacteria while the antibiotic targets phage-resistant (antibiotic-sensitive) mutants^7,11^. Critically, in this model we assume that immune effector cells can target both bacterial strains. As we show, combination therapy successfully clears infections insofar as immune responses are active. Our proof-of-principle systems-level model highlights the role of immune responses in developing and assessing the effectiveness of phage-based therapeutics for treatment of MDR pathogens, particularly MDR *P.a*. that exhibit evolutionary trade-offs.

## II. COMBINATION THERAPY MODEL

We propose a combination therapy model consisting of a system of nonlinear, ordinary differential equations representing the interactions among bacteria, phage, antibiotics, and the innate immune system (see Figure 1). Two strains of bacteria are included, one of which is phage-sensitive (*B*_*P*_) and the other of which is antibiotic-sensitive (*B*_*A*_). The strains *B*_*P*_ and *B*_*A*_ reproduce given limitation by a carrying capacity. *B*_*P*_ is infected and lysed by phage (*P*) but resists the antibiotic, while *B*_*A*_ population is killed by the antibiotic but is resistant to phage (for an *in vitro* model of bacteriophage therapy with fully susceptible and resistant types see^17^). We do not consider double-resistant mutants in our model due to the evolutionary trade-off between resistance against phage and antibiotics observed for *P.a.*^7^. Phage replicate inside the host *B*_*P*_ and decay in the environment. The antibiotic is administered at a constant concentration, then it is metabolized and removed at a fixed rate. The population dynamics is governed by the set of equations:

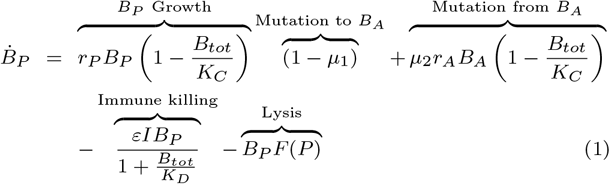

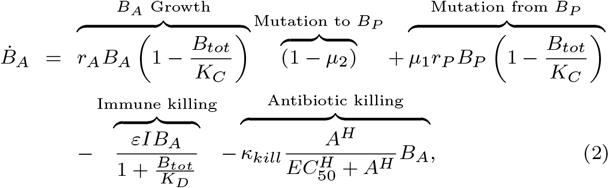

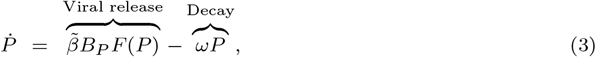

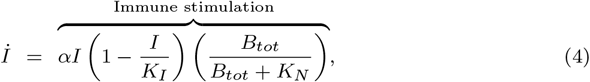

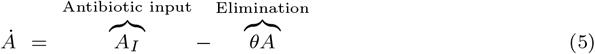

**FIG. 1:**
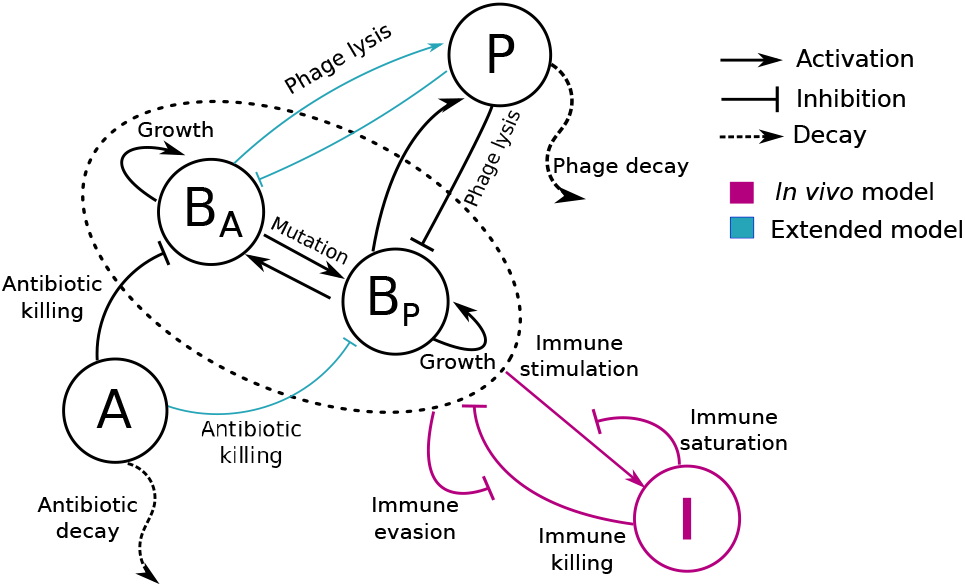
Schematic of the phage-antibiotic combination therapy model. Antibiotic-sensitive bacteria (*B*_*A*_) and phage-sensitive bacteria (*B*_*P*_) are targeted by antibiotic (A) and the phage (P), respectively. Host innate immune response interactions (pink arrows) are included in the *in vivo* model. Innate immunity (I) is activated by the presence of bacteria and attacks both bacterial strains. Furthermore, in model versions accounting for partial resistance (blue arrows), *B*_*A*_ and *B*_*P*_ are targeted by both antibiotic and phage but in quantitatively different levels.

In this model, phage-sensitive bacteria grow at a maximum rate *r*_*P*_, while antibiotic-sensitive bacteria (*B*_*A*_) grow at a maximum rate *r*_*A*_. The total bacterial density, *B*_*tot*_ = *B*_*P*_ + *B*_*A*_, is limited by the carrying capacity *K*_*C*_. Phage infect and lyse *B*_*P*_ bacteria at a rate *F*(*P*). Antibiotic killing is approximated by a Hill function with the nonlinear coefficient (*H*)^18–21^. The maximum antibiotic killing rate is *κ*_*kill*_, while *EC*_50_ is the concentration of the antibiotic, here considered as ciprofloxacin, at which the antibiotic effect is half as the maximum. Phage *P* replicate with a burst size 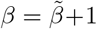 and decay at a rate *ω*. We assume that antibiotic dynamics are relatively fast, and use a quasi-steady state approximation of *A*^∗^ = *A*_*I*_/*θ*.

When simulating an *in vivo* scenario, the host innate immune response, *I*, is activated by the presence of bacteria and increases with a maximum rate *α*. *K*_*N*_ is a half-saturation constant, i.e., the bacterial density at which the growth rate of the immune response is half its maximum. Bacteria grow and are killed by the innate immunity with a maximum killing rate *ε*. However, at high bacterial concentration bacteria can evade the immune response and reduce the immune killing efficiency^5,16^.

Our model uses an implicit representation of spatial dynamics through different functional forms of phage-bacteria interactions (*F*(*P*)). As such, we do not explicitly model the spatial dynamics of individual components. The model considers three modalities of phage infection, *F*(*P*): linear, heterogeneous mixing^5,22^, and phage saturation^5^. The linear phage infection modality assumes a well-mixed environment, where phage easily encounter and infect bacteria, so the infection rate *F*(*P*) = *ϕP* is proportional to the phage density, where *ϕ* is the linear adsorption rate. The heterogeneous mixing model accounts for spatial heterogeneity, 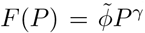. Where 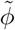 is the nonlinear adsorption rate and *γ* < 1 the power-law exponent. The third modality assumes that at high phage density multiple phage particles adsorb to a single bacterium so phage infection follows a saturating Hill function, 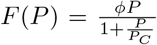. Here, *ϕ* is the adsorption rate and *P*_*C*_ is the phage density at which the infection rate is half-saturated.

Note that in later stages, we consider an ‘extended’ combination therapy model (Figure 1 (blue arrows)) in which bacterial strains are sensitive to both phage and antibiotic in quantitatively distinct levels. The full set of equations for this extension are found in the Supplementary Materials. In addition, a full description of parameter choices are described in the Methods.

## III. RESULTS

### A. Differential outcomes of single phage therapy

We begin by exploring the dynamics arising from adding a single phage type at a density of 7.4 × 10^8^ PFU/g two hours after infections caused by either phage-sensitive or phage-resistant bacteria (Fig. 2). When the infection is caused by a phage-sensitive bacteria (*B*_*P*_ = 7.4 × 10^7^ CFU/g), phage lysis reduced *B*_*P*_ density to the point where the immune response alone could control this bacterial population. Despite the emergence of phage-resistant mutants (*B*_*A*_), total bacterial population remained low and the innate immunity effectively controlled the infection. On the other hand, when the infection was caused by phage-resistant mutants (*B*_*A*_ = 7.4 × 10^7^ CFU/g), the phage could not target *B*_*A*_ so the bacterial population grew unimpeded. The immune response was overwhelmed by the rapid growth of *B*_*A*_ which then reached a density of ~10^10^ CFU/g after 12h (Fig. 2 bottom), leading to a persistent infection despite an activated immune response (similar to the outcomes described in^16^).

**FIG. 2:**
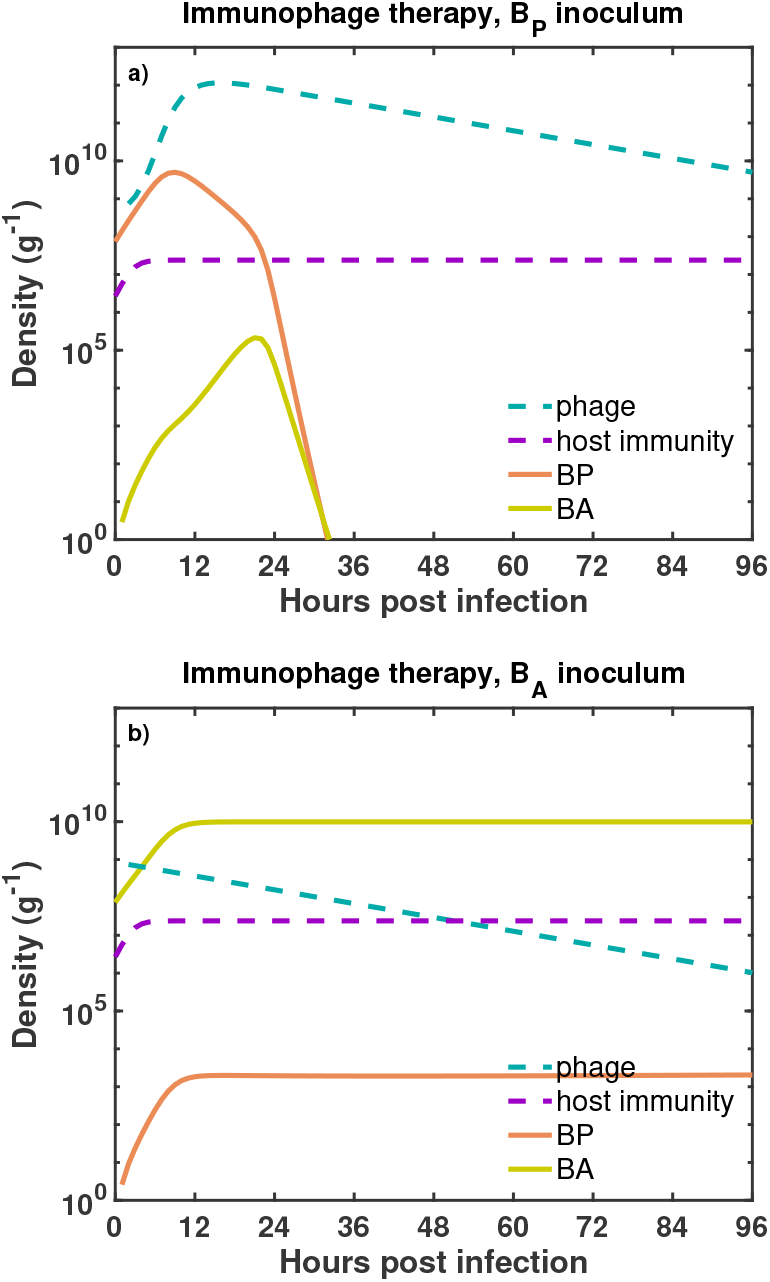
Dynamics of the immunophage therapy model against two different bacterial inoculum. We simulate the phage therapy model developed by^16^ against two infection settings. In the first infection setting (a), a phage-sensitive bacterial inoculum, *B*_*P*_ (orange solid line), is challenged with phage (blue dashed line) inside an immunocompetent host. In the second scenario (b), antibiotic-sensitive bacteria, *B*_*A*_ (green solid line), are challenged with phage in presence of an active immune response (purple dashed line). The initial bacterial density and the initial phage density are, *B*_0_ = 7.4 × 10^7^ CFU/g and *P*_0_ = 7.4 × 10^8^ PFU/g, respectively. The growth rates of *B*_*P*_ and *B*_*A*_ are, *r*_*P*_ = 0.75*h*^−1^ and *r*_*A*_ = 0.67*h*^−1^, respectively. Simulation run is 96 hours with phage being administered 2 hours after the infection. The bacterial carrying capacity is, *K*_*C*_ = 10 CFU/g.

This initial analysis illustrates how therapeutic outcomes given application of a single phage type may be strongly dependent on the initial bacterial inoculum. As expected, single phage therapy fails to clear the infection when the bacterial inoculum is mistargeted (Fig. 2 bottom). In the next section we evaluate infection dynamics in response to the combined application of phage and antibiotics – similar to that in multiple *in vitro* and *in vivo* studies of phage-antibiotic treatment of MDR *P.a.*.^7,11^.

### B. Phage-antibiotic therapy treatment dynamics in immunocompetent hosts

We simulated the combined effects of phage (7.4 × 10^8^ PFU/g) and antibiotics (assuming 2.5×MIC of ciprofloxacin for the *B*_*A*_ strain) in two different infection settings. First, when an immunocompetent host was infected with phage-sensitive bacteria, the infection was cleared before ~36 hours due to the combined killing effect of phage, antibiotic, and innate immunity. The dominant bacterial population, *B*_*P*_, was targeted by the phage while the antibiotic targeted *B*_*A*_. The combined effects of phage and antibiotic reduced total bacterial density to the point where innate immunity eliminate the bacterial infection. Second, when the host was infected with antibiotic-sensitive bacteria, the pathogen was cleared (before ~12 hours) due to the combined effect of phage, antibiotic, and innate immunity. The antibiotic facilitated the decrease of *B*_*A*_ while phage maintained *B*_*P*_concentration low, easing the innate immunity control over the infection. The resulting infection clearance in the phage-resistant case (Fig. 3 bottom) stands in stark contrast to the previous outcome of the single phage therapy model (Fig. 2 bottom). Overall, the results suggest that a curative outcome is possible when phage are combined with antibiotics in an immunocompetent host – even when the phage is initially mistargeted to the dominant bacterial strain. However what remains unclear is the extent to which successful treatment is driven by phage and antibiotics alone or, in part, because of the synergistic interactions with the innate immune response.

**FIG. 3:**
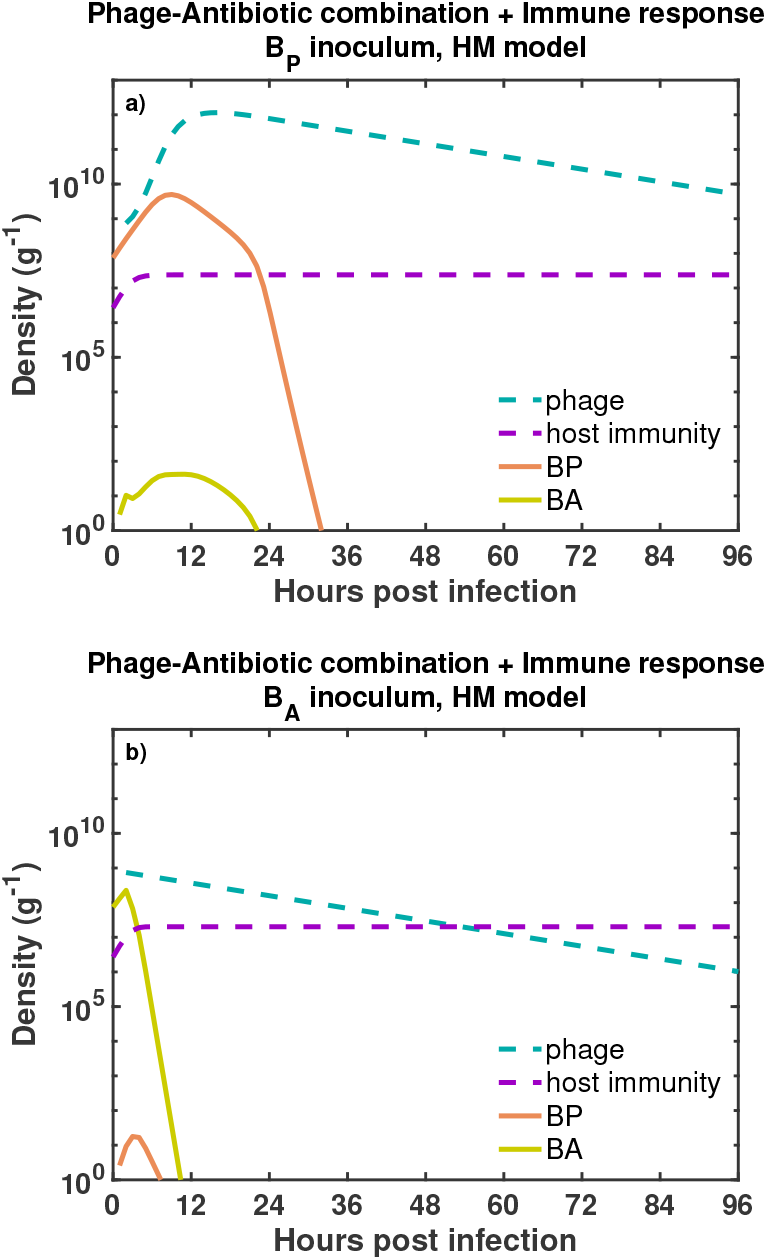
Outcomes of the phage-antibiotic combination therapy model for two different infection settings. We simulate the combined effects of phage and antibiotics in an immunocompetent host infected with phage-sensitive bacteria (a), *B*_*P*_ (orange solid line). In (b), the host is infected with antibiotic-sensitive bacteria, *B*_*A*_ (green solid line). The dynamics of the phage (blue dashed line) and innate immunity (purple dashed line) are shown for each infection setting. Initial bacterial density and phage density are, *B*_0_ = 7.4 × 10^7^ CFU/g and *P*_0_ = 7.4 × 10^8^ PFU/g, respectively. The simulation run is 96 hours (4 days). Antibiotic and phage are administered 2 hours after the beginning infection. Ciprofloxacin is maintained at a constant concentration of 0.0350 *µ*g/ml during the simulation. The carrying capacity of the bacteria is *K*_*C*_ = 10^10^ CFU/g.

### C. Phage-antibiotic combination therapy requires innate immunity to robustly clear the pathogen

In this section we assess the dependency of combination therapy on the immune response. To do so, we evaluate the combination therapy while setting *I* = 0. This is meant to mimic conditions of severe immunodeficiency. In order to further assess outcomes, we also consider multiple functional forms for phage-bacteria interactions - including the phage-saturation, heterogeneous mixing, and linear infection models (see Methods for more details).

First, when a phage-sensitive bacterial inoculum was challenged with the combination therapy, the pathogen persisted in two of three infection models. Bacteria persist in the HM (Fig. 4 top-left) and PS (Fig. 4 upper-middle) models, while the combination of phage and antibiotic successfully eliminates the bacterial population in the LI model (Fig. 4 top-right). Although the combination of phage and antibiotic did not eliminate the bacterial population in the HM and PS models, the combination strategy still reduced the bacterial concentration relative to the carrying capacity (*K*_*C*_ = 10^10^ CFU/g).

**FIG. 4:**
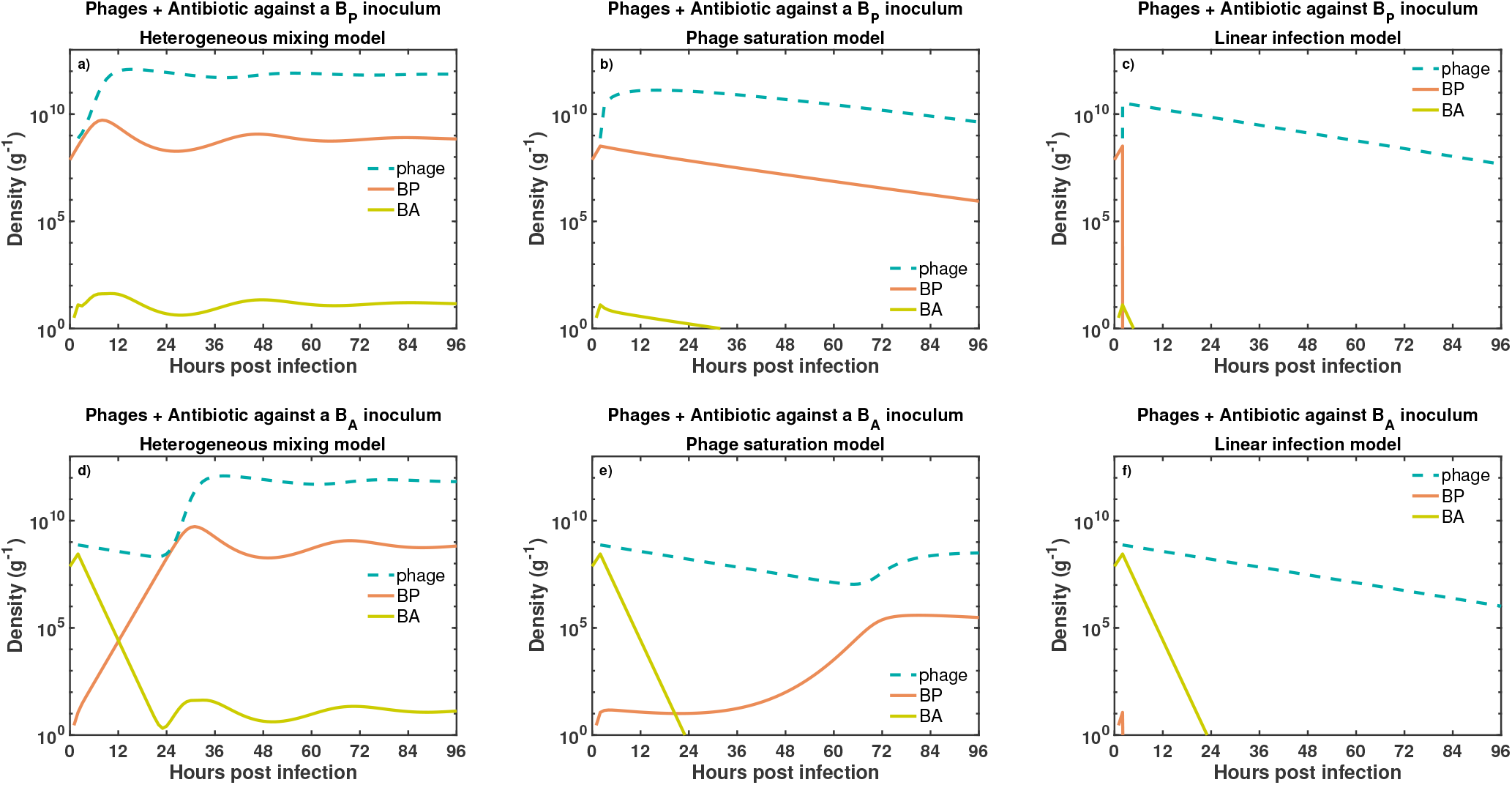
Bacterial dynamics given joint exposure to phage and antibiotic. We simulate bacterial growth for 96 hours in exposure to phage (blue dashed line) and antibiotic (data not shown) added two hours after the beginning of the inoculation. The combination of phage and antibiotic is tested against two different bacterial inoculum. The first inoculum consisted of exclusively phage-sensitive bacteria (a-c), *B*_*P*_ (orange solid line). The second inoculum consisted of antibiotic-sensitive bacteria (d-f), *B*_*A*_ (green solid line). Additionally, we test three different models of phage infection, heterogeneous mixing (a, d), phage saturation (b, e) and linear infection models (c, f). The initial bacterial density and phage density are, *B*_0_ = 7.4 × 10^7^ CFU/g and *P*_0_ = 7.4 × 10^8^ PFU/g, respectively. Ciprofloxacin is maintained at a constant concentration of 2.5×MIC (i.e., 0.0350 *µg/ml*) during the simulations.

Second, when an antibiotic-sensitive bacterial inoculum was challenged with phage and antibiotic, bacteria persisted in two of three infection models, similar to the previous phage-sensitive case. Bacteria persist in the HM (Fig. 4 bottom-left) and PS (Fig. 4 lower-middle) models, while bacterial population is eliminated in the LI model (Fig. 4 bottom-right). Inclusion of antibiotics facilitated a decrease in *B*_*A*_ and the spread of *B*_*P*_leading to coexistence between bacteria and phage. Furthermore, the elimination of bacteria in the LI model took longer (~24 hours) compared to the previous phage-sensitive case.

The outcomes of the combination therapy model suggest that, in absence of innate immunity, infection clearance is not achieved in two of three phage infection models. Pathogen clearance is only achieved in the linear infection case, that is, when we assume a well-mixed environment. On the other hand, when we assume spatial heterogeneity or phage saturation, a coexistence state between phage and bacteria arises from the tripartite dynamics between phage, bacteria, and antibiotic. Such coexistence state is inconsistent with the expected antimicrobial effect of the combination therapy^7^ – and points to a potentially unrealized role of the immune response in the effectiveness of phage-antibiotic combination therapy.

### D. Outcomes of the combination therapy model are robust to the bacterial composition of the inoculum and the concentration of antibiotic

Thus far we have simulated two extreme infection inoculum scenarios involving exclusively phage-sensitive bacteria or exclusively antibiotic-sensitive bacteria. Next, we consider the effects of combination therapy on mixed bacterial inoculum containing both *B*_*P*_ and *B*_*A*_. To do so, we performed a robustness analysis of four (*in silico*) therapy models, i.e., antibiotic-only, antibiotic-innate immunity, phage-antibiotic, and phage-antibiotic combination in presence of innate immunity. For each model, we varied the concentration of the antibiotic and the bacterial composition of the inoculum. Outcomes from the different therapeutics are consistent with previous results obtained using a fixed set of initial conditions (Table I). We find that model outcomes are robust to variations in the initial conditions (i.e., inoculum composition and concentration of ciprofloxacin).

First, we evaluated the killing effect of the antibiotic against mixed bacterial inoculum. We find that the pathogen persisted (~ 10^10^ CFU/g) for all different inoculum and concentrations of antibiotic. The antibiotic targeted *B*_*A*_ while *B*_*P*_ grew unimpeded in the absence of phage, such that *B*_*P*_ predominated after 96 hrs. In contrast, antibiotics and innate immunity (Fig. 5 top-right) could eliminate bacterial inoculum with high percentages of antibiotic-sensitive bacteria (*>* 90% of B_A_). During this scenario, the low percentages of *B*_*P*_coupled to the antibiotic killing of *B*_*A*_ facilitated the immune clearance of the infection. Furthermore, pathogen clearance was observed even for sub-inhibitory concentrations of ciprofloxacin. As is apparent, the antibiotic on its own cannot clear the infection, and therapeutic outcomes are only modestly improved in a narrow region of inoculum space.

**FIG. 5:**
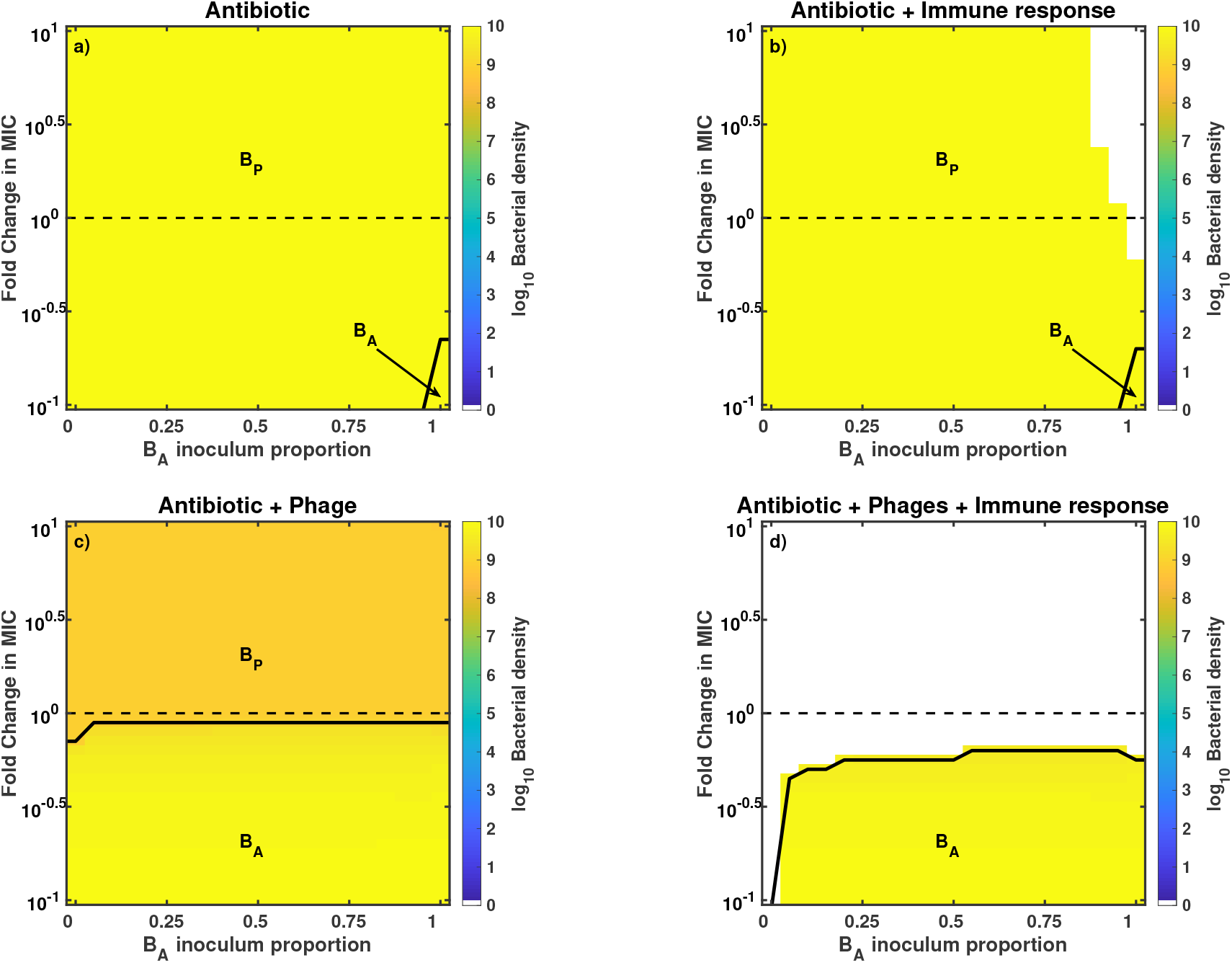
Outcomes of the robustness analysis for different antimicrobial strategies. We simulate the exposure of bacteria to different antimicrobial strategies, such as antibiotic-only (a), antibiotic + innate immunity (b), phage + antibiotic (c), and phage-antibiotic combination in presence of innate immunity (d). The heatmaps show the bacterial density at 96 hours post infection. Colored regions represent bacteria persistence (e.g., orange areas ~ 10^9^ CFU/g and bright yellow areas ~ 10^10^ CFU/g) while the white regions represent pathogen clearance. We vary the concentration of ciprofloxacin (MIC = 0.014 *µ*g/ml), ranging from 0.1 MIC (0.0014 *µ*g/ml) to 10 MIC (0.14 *µ*g/ml), and the bacterial composition of the inoculum ranging from 100% phage-sensitive bacteria (0% *B*_*A*_) to 100% antibiotic-sensitive bacteria (100% *B*_*A*_). Initial bacterial density and phage density (c-d) are, *B*_0_ = 7.4 × 10^7^ CFU/g and *P*_0_ = 7.4 × 10^8^ PFU/g, respectively. Phage and antibiotic are administered two hours after the beginning of the infection.

**TABLE I:** Summary of therapeutic outcomes given a combination of antibiotics (*A*), phage (*P*), and immunity (*I*). The presence or absence of different antimicrobial agents is represented with 1 or 0, respectively.

| Treatment |  |  | Outcome |
| --- | --- | --- | --- |
| A | P | I |  |
| 1 | 0 | 0 | Infection via BP proliferation |
| 1 | 0 | 1 | Infection via BP proliferation |
| 1 | 1 | 0 | Infection via BP coexistence with phage |
| 1 | 1 | 1 | Curative |

Second, we assessed the effects of combining antibiotics with phage against mixed bacterial inoculum. The phage-antibiotic combination strategy failed to clear the infection for all combinations of initial conditions – consistent with the infection scenarios of the prior section. Nonetheless, bacterial concentration was ~10 times smaller due to phage killing (orange area in Fig. 5 bottom-left) compared to the bacterial concentration from the antibiotic-only therapy (bright yellow area of Fig. 5 top-left). After 96 hours of combined treatment, the phage-sensitive population was predominant at above MIC levels while antibiotic-sensitive bacteria populated the sub-MIC levels (Fig. S1 and Fig. S2 show the effects of different antibiotic levels on the bacterial dynamics). In contrast, a robust pathogen clearance was achieved when the phage-antibiotic combination strategy was supplemented with active innate immunity (Fig. 5 bottom-right). Note that, even partially effective immune responses can still be sufficient to achieve infection clearance (Fig. S3). Overall, the synergistic interactions between phage, antibiotic, and innate immunity led to clearance of the infection for the majority of initial conditions. The clearance region even spanned sub-inhibitory concentrations of ciprofloxacin.

We performed a further exploratory analysis of the combined therapy. We studied the effects of delay times on the application of the combined strategy, showing that therapeutic action is robust to delay times and fails irrespective of delay time when the immune system is compromised (Text S1, Fig. S4). We also performed a parameter sensitivity analysis (Text S1), showing that the combined strategy, when supplemented with the host immune response, is effective for a wide range of parameters (Fig. S5).

Finally, we note that these results derived from analysis of dynamics arising among extreme phenotypes. In reality, phage-sensitive strains may retain some sensitivity to antibiotics and antibiotic-sensitive strains can be infected at reduced levels by phage^7,23,24^. Hence, we repeated the robustness analysis, using an extended model that incorporates quantitatively different levels of phage-infectivity and antibiotic-sensitivity of both strains (see Supplementary Text S2). Partial resistance model outcomes are qualitatively consistent with previous outcomes of the extreme resistance model (contrast Fig. S6 with Fig. 5). Moreover, the bacterial dynamics of the partial resistance model are qualitatively similar to the dynamics arising among extreme phenotypes (contrast Fig. 3 with Fig. S7-bottom). Overall, our model analysis suggests that robust, curative success of phage-antibiotic combination therapy could be driven, in part, by a largely unrealized synergy with the immune response.

## IV. DISCUSSION

We have developed a combination therapy model that combines phage and antibiotics against a mixed-strain infection of *Pseudomonas aeruginosa*. The model suggests that infection clearance arises from nonlinear synergistic interactions between phage, antibiotic, and innate immunity. Moreover, the infection clearance shows robustness to variations in the concentration of antibiotic, delays in the administration of the combined therapy, the bacterial composition of the inoculum, and model assumptions. In contrast, when innate immunity responses are removed (or severely reduced), then phage-antibiotic combination therapy is predicted to fail to eliminate the infection. This suggests that combined therapy may depend critically on immune response for resolving bacterial infections.

The in sillco findings are consistent with qualitative, experimental outcomes *in vitro* and *in vivo*. For example, one of our main results states that phage-antibiotic combined therapy has a greater antimicrobial effect than single phage or antibiotic therapies, this is consistent with several *in vitro* settings that show a greater bacterial density reduction for combined rather than single therapies^25–28^. Moreover, additional studies explore the use of sub-lethal concentrations of antibiotics otherwise insufficient for controlling bacterial growth but efficient when combined with phage against diverse bacterial populations^25,26,28,29^. These findings are consistent with our *in silico* outcomes where pathogen clearance is observed at sub-MIC antibiotic levels in the combined therapy framework. Further work to compare model-based predictions to experiments will require moving beyond outcomes to high-resolution temporal data.

In connecting models to experiment, it is important to consider extending the model framework to a spatially explicit context. Spatial structure can be relevant therapeutically. For example, during chronic infections spatially organized bacterial aggregates of *P.a.* protect themselves against phage killing by producing exopolysaccharides^30^. Furthermore, modelling efforts have shown that spatial structure affects the therapeutic success of phage therapy^31^ and phage-antibiotic combination therapy^32^. For example, structured environments limit phage dispersion and amplification, promoting bacterial survival and resistance acquisition^31,32^. Moreover, the heterogeneous distribution of antibiotic creates spatial refuges (of low or null antimicrobial presence) where bacteria survive and resistant mutants arise^32^. The current model also neglects the complex features of immune response termination^33^ and interactions with commensal microbes^34^, both priority areas for future work.

In conclusion, the phage-antibiotic combination therapy model developed here describes efforts to explore how host immunity modulates infection outcomes. As we have shown: immune clearance of pathogens may lie at the core of the curative success of combination treatments. If so, this additional synergy may help to resolve the resistance problem and also guide use of sub-MIC concentrations of antibiotics. Besides reducing toxic side effects associated with high concentrations of antibiotics, sub-MIC concentrations can improve phage infectivity through morphological changes of the bacterial cell^9,35,36^ or by not interfering with the phage replication cycle^25,26^. When combined in an immunocompetent context, we find that phage-antibiotic combination therapy is robust to quantitatively and qualitatively distinct resistance profiles. These findings reinforce findings that phage and antibiotics can be used to treat a certain class of MDR *P.a.* pathogens in patients^11,37^. Model results also highlight the role of the immune response in realizing curative success - which will be relevant to expanding combination theory for a range of clinical applications.

## V. METHODS

### A. Model simulation

The numerical integration of the combination therapy model is carried out using ODE45 in MATLAB. We obtain the temporal dynamics of two bacterial strains, phage, antibiotic, and innate immune response. Moreover, we set an extinction threshold of 1 g^−1^; hence, when *B*_*P*_ or *B*_*A*_ densities are ≤ 1 CFU/g at any time during the simulation, we set their densities to 0 CFU/g. We run all the simulations for 96 hours (4 days).

### B. Robustness analysis

We perform a robustness analysis of the phage-antibiotic combination therapy model by varying its initial conditions. We vary the concentration of antibiotic from sub-MIC concentrations (0.1 MIC) to above MIC concentrations (10 MIC), using the MIC of ciprofloxacin (0.014*µ*g/ml) for PAPS phage-resistant strain as a reference^11^. Moreover, we vary the bacterial composition of the inoculum by increasing the bacterial density of one strain (e.g., *B*_*A*_) by 5% and decreasing the density of the other by 5%. Then we select a pair of initial conditions and run the model 96 hrs. Finally, we calculate total bacterial density, *B*_*total*_ = *B*_*A*_ + *B*_*P*_.

### C. Parameter Estimation

The parameter values used in the simulations of the combination therapy model are shown in Table II–III. Most of the parameter estimation was carried out in previous work (see *Parameter Estimation* section in Ref.^5^), supplemented by parameters associated with functions describing the pharmacodynamics and pharmacokinetics of ciprofloxacin^18,38^.

**TABLE II:**
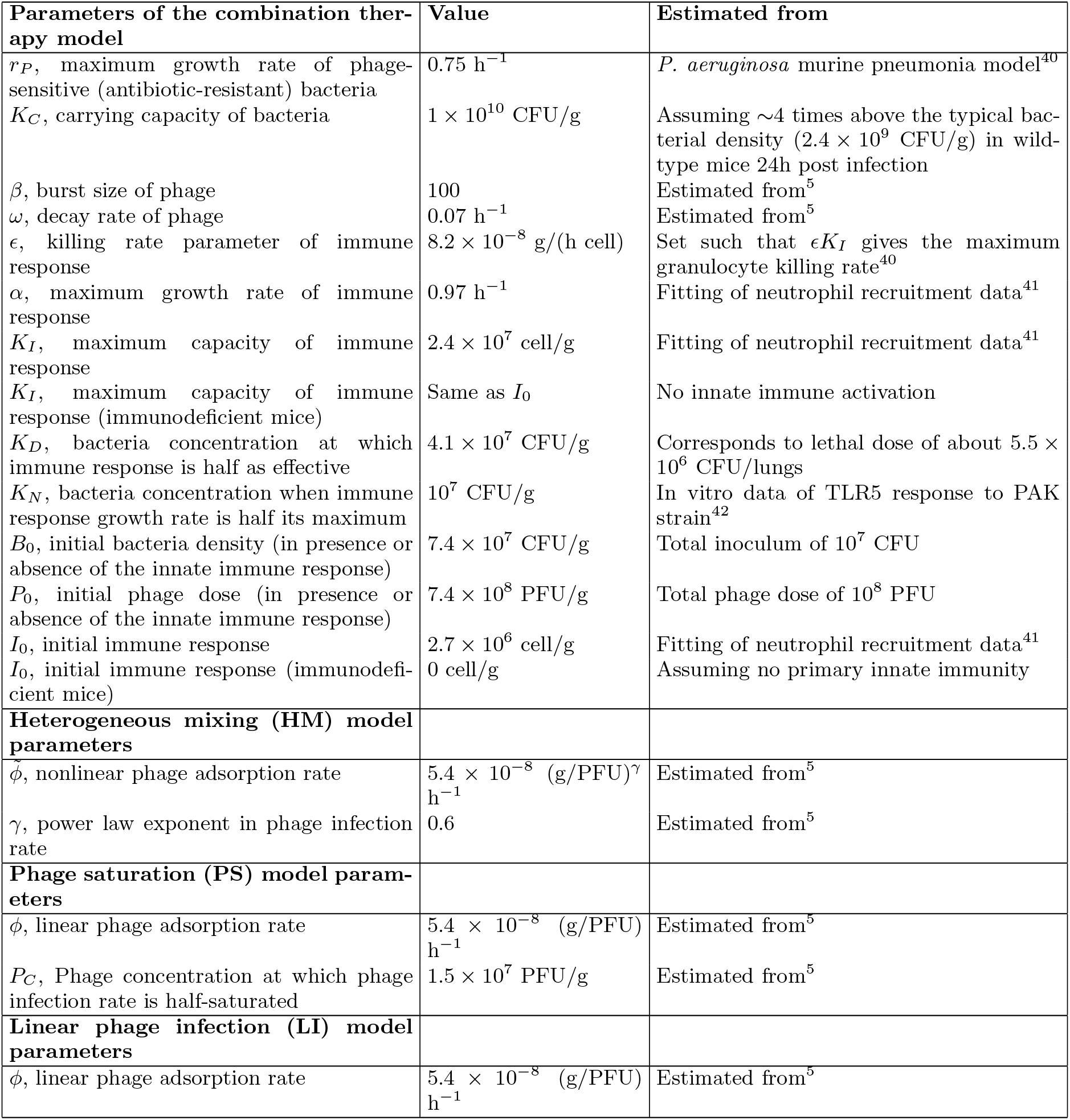
Microbiology and phage-associated parameter values.

**TABLE III:** Additional parameter values associated with the effects of antibiotics.

| **Antibiotic (ciprofloxacin) parameters** |  |  |
| --- | --- | --- |
| $\kappa_{kill}$, maximum antibiotic killing rate | 18.5 h<sup>-1</sup> | Fitting an Emax model to antibiotic kill curves<sup>18</sup> |
| $EC_{50}$, concentration of antibiotic at which the killing rate is half its maximum | 0.3697 $\mu g/ml$ | Calculated using the MIC of ciprofloxacin for the phage-resistant PAPS strain<sup>7</sup> |
| $EC_{BP}$, concentration of antibiotic at which the killing rate is half its maximum | 4.070 $\mu g/ml$ | Calculated using the MIC of ciprofloxacin for the phage-sensitive PAPS strain<sup>7</sup> |
| $H$, hill coefficient | 1 | From<sup>18</sup> |
| $MIC$, Minimum inhibitory concentration of ciprofloxacin for *P.a.* PAPS phage-resistant strain | 0.014 $\mu g/ml$ | From<sup>7</sup> |
| $MIC$, Minimum inhibitory concentration of ciprofloxacin for *P.a.* PAPS phage-sensitive strain | 0.172 $\mu g/ml$ | From<sup>7</sup> |
| $\theta$, antibiotic elimination rate from serum samples | 0.53 h<sup>-1</sup> | Estimated from antibiotic concentration vs time curves; concentration of ciprofloxacin was measured in serum samples of *P.a.*-infected mice<sup>38</sup> |
| **Antibiotic-sensitive bacteria parameters** |  |  |
| $\mu_1$, probability of emergence of antibiotic-sensitive (phage-resistant) mutants per cellular division | $2.85 \times 10^{-8}$ | Estimated from experimental measurements<sup>39</sup> |
| $\mu_2$, probability of emergence of phage-sensitive (antibiotic-resistant) mutants per cellular division | $2.85 \times 10^{-8}$ | Approximated to the estimates from<sup>39</sup> |
| $r_A$, maximum growth rate of antibiotic-sensitive (phage-resistant) bacteria | 0.675 h<sup>-1</sup> | 10% trade-off between resistance against phage and growth rate<sup>43</sup> |

The pharmacodynamics of ciprofloxacin (CP) is described by the following Emax model^18^,

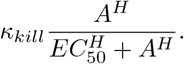

Where *κ*_*kill*_ represents the maximum killing rate of the antibiotic, *EC*_50_ is the antibiotic concentration at which the antibiotic killing rate is half its maximum, and *H* is a Hill coefficient. The values of the parameters are obtained using *in vitro* growth curves of *P.a.* at different concentrations of CP^18^. The elimination rate of the antibiotic, *θ*, is estimated from levels of clearance of CP from serum samples of mice infected with *P.a.*^38^. The *EC*_50_ parameter value is adjusted in our model to consider the MIC of CP for PAPS reference strains^7^.

The probabilities of producing a mutant strain per cell division, *µ*_1_ and *µ*_2_, are obtained from Ref.^39^. Where *µ*_1_ is the probability of producing a phage-resistant mutant (antibiotic-sensitive) per cell division and *µ*_2_ is the probability of generating a phage-sensitive (antibiotic-resistant) mutant per cell division.

To account for partially resistant strains, we extend our combination therapy model (Equations S21-S25) and include the parameters, 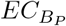 and *δ*_*P*_. 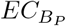 is the half-saturation constant of the antibiotic killing function and modulates the level of antibiotic resistance for *B*_*P*_. The parameter was calculated based on the MIC of CP for PAPS phage-sensitive strain^7^. Moreover, for modulating the level of resistance to phage infection (*F*(*P*)) of *B*_*A*_ we use the parameter *δ*_*P*_ < 1 (see Table IV).

**TABLE IV:** Phage adsorption rate of phage-sensitive and phage-resistant *Pseudomonas aeruginosa* strains.

| Phage adsorption rate of phage-sensitive strain (%) | Phage adsorption rate of phage-resistant strain (%) | Fold decreased phage adsorption rate | Ref |
| --- | --- | --- | --- |
| 31.3% | 9.6% | 3.2 | 23 |
| 73.7% | 66.5% to 2.4% | $\sim 1.1$ to $\sim 30$ | 44 |
| 87.2% | 42.8% | 2 | 44 |
| 92.7% | 55.3% and 42.8% | 1.6 and 2 | 24 |

## Supplemental Material

**Text S1**. Robustness and sensitivity analysis of the combination therapy model.

**Text S2**. Partial resistance model, an extension of the combination therapy model.

Fig S1. Bacterial dynamics given exposure to low levels of antibiotics.

Fig S2. Accounting for variations in the concentration of antibiotic, from sub-MIC to MIC levels.

Fig S3. Bacterial density after 96 h of combined treatment with intermediate immune response levels.

Fig S4. Time delays in the application of the combined treatment.

Fig S5. Parameter sensitivity analysis results.

Fig S6. Robustness analysis of the partial resistance model for different antimicrobial strategies.

Fig S7. Time series of the partial resistance model in presence and absence of host immune response.

## Acknowledgements

The authors thank S. Brown, J. Gurney, and K. Kortright for discussions. The work was supported by a grant from the Army Research Office W911NF-14-1-0402 (to JSW), a grant from the National Science Foundation NSF PoLS 1806606 (to JSW), and a pilot award from the Cystic Fibrosis Foundation (to BKC and PET).

## Supplemental Text S1: Robustness and sensitivity analysis of the combination therapy model

We performed a robustness analysis of the combined therapy model, where we varied the initial conditions such as concentration of antibiotics and composition of the bacterial inoculum (Fig. 5). Here, we analyze the effects of such variations on the bacterial dynamics. For example, the model predicts that when low or null concentrations of antibiotic are used against inoculum with non-trivial levels of antibiotic-sensitive bacteria *B*_*A*_, the infection persists and *B*_*A*_ strain dominates after 96 hours (Fig. S1). On the other hand, infection clearance is obtained once we increase the concentration of the antibiotic, e.g., close to the MIC (Fig. S2-A), indicating that higher concentrations are needed to eliminate the infection for inoculum with non-trivial levels of *B*_*A*_.

We further explore the claim that a sufficient level of immune response is needed to achieve a robust infection clearance. Hence, we varied the levels of innate immune activation from 20% to 100% for the A + P + I regime and simulated the combined therapy model for 96 hours. We predict that at least 60% of immune activation is needed to achieve a robust pathogen clearance (Fig. S3). We also explored the effects of time delays on the combination therapy. Phage and antibiotics were applied simultaneously 2 to 10 hours after the beginning of the infection, obtaining qualitatively consistent outcomes for this range of time-delays (Fig. S4).

Finally, we performed a parameter sensitivity analysis to evaluate the robustness of our model outcomes to parameter variations. Because parameters are adapted from experimental data whenever possible, we allow parameters to vary up to 10% of their original values. Then, we focused on two therapeutic regimes, A + P + I and A + P, and asked what fraction of the domain range in MIC and *B*_*A*_ proportion led to complete elimination (white regions on A + P + I heatmaps); this value is 62% for A + P + I and 0% for A + P when the reference parameter set, *θ*_*ref*_, is used. We iterate this process 1000 times using different perturbed parameter sets, *θ*_*per*_, and obtain a distribution of the percentage of complete elimination for the two therapeutic regimes. We found that ~ 2/3 of the total runs led to a fraction of 50% or more complete elimination (Fig. S5) for the A + P + I regime while no elimination was observed for the A + P regime in 1000 iterations. Overall, the sensitivity analysis supports the claim that joint synergistic effects of phage, antibiotics, and the immune response occur for many (rather than a particular choice of) parameter sets.

## Supplemental Text S2: Partial resistance model, an extension of the combination therapy model

The extended *in vivo* combination therapy model describes a system where two bacterial strains partially sensitive to both, phage and antibiotic, interact with an active host immune response:

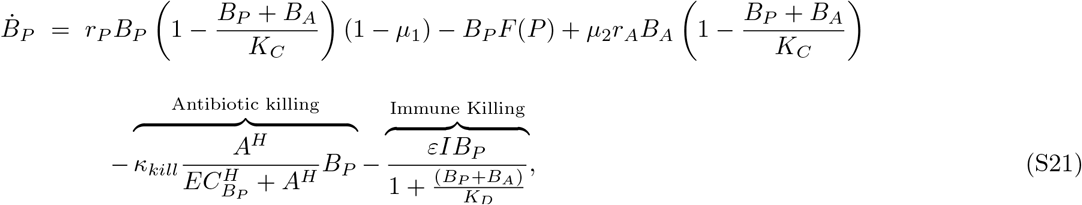

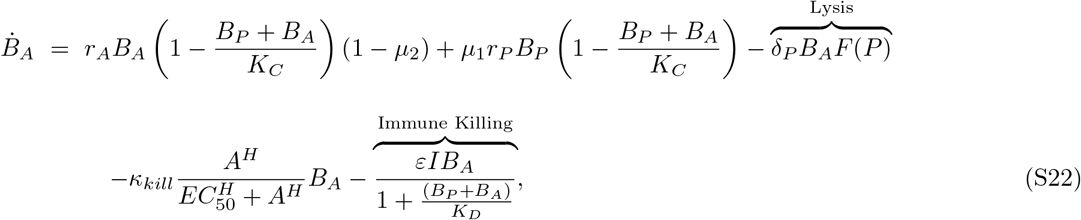

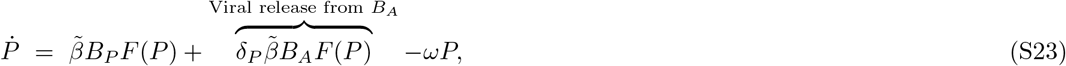

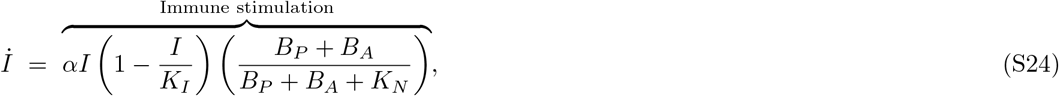

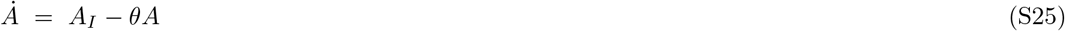

In this model, the parameter 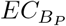 modulates the level of antibiotic resistance of the phage-sensitive bacteria, *B*_*P*_ (Equation S21) and 0 ≤ *δ*_*P*_ ≤ 1 modulates the level of phage-resistance of *B*_*A*_ (Equation S22). Finally, we add a new viral release term in the phage dynamics that accounts for phage infection of antibiotic-sensitive bacteria (Equation S23).

We simulate the partial resistance model covering immunodeficient and immunocompetent states (Fig. S7 top and bottom, respectively). We applied the combined treatment against two different infection settings, in the first setting the inoculum consisted of exclusively phage-sensitive bacteria and the second inoculum was composed of exclusively antibiotic-sensitive bacteria. Our simulations predict that the combined treatment may fail to clear the infection under an immunodeficient state regardless of the bacterial composition of the inoculum. This result is consonant with our previous finding (compare Fig. S7-top with Fig. 4). On the other hand, the incorporation of an active innate immunity may facilitate the elimination of the infection regardless of the bacterial genotype, i.e., phage-sensitive (Fig. S7, bottom-left) or antibiotic-sensitive (Fig. S7, bottom-right).

The combined effect between phage, antibiotic, and innate immune response leads to a synergistic infection clearance (Fig. S6, bottom-right). Our results suggest that even for bacterial strains that remain partially sensitive to both phage and antibiotics, the presence of the host innate immunity is still necessary to clear the infection. Hence, the outcomes of the combination therapy model are robust to model extensions that account for partially resistant strains.

---

**FIG. S1:**
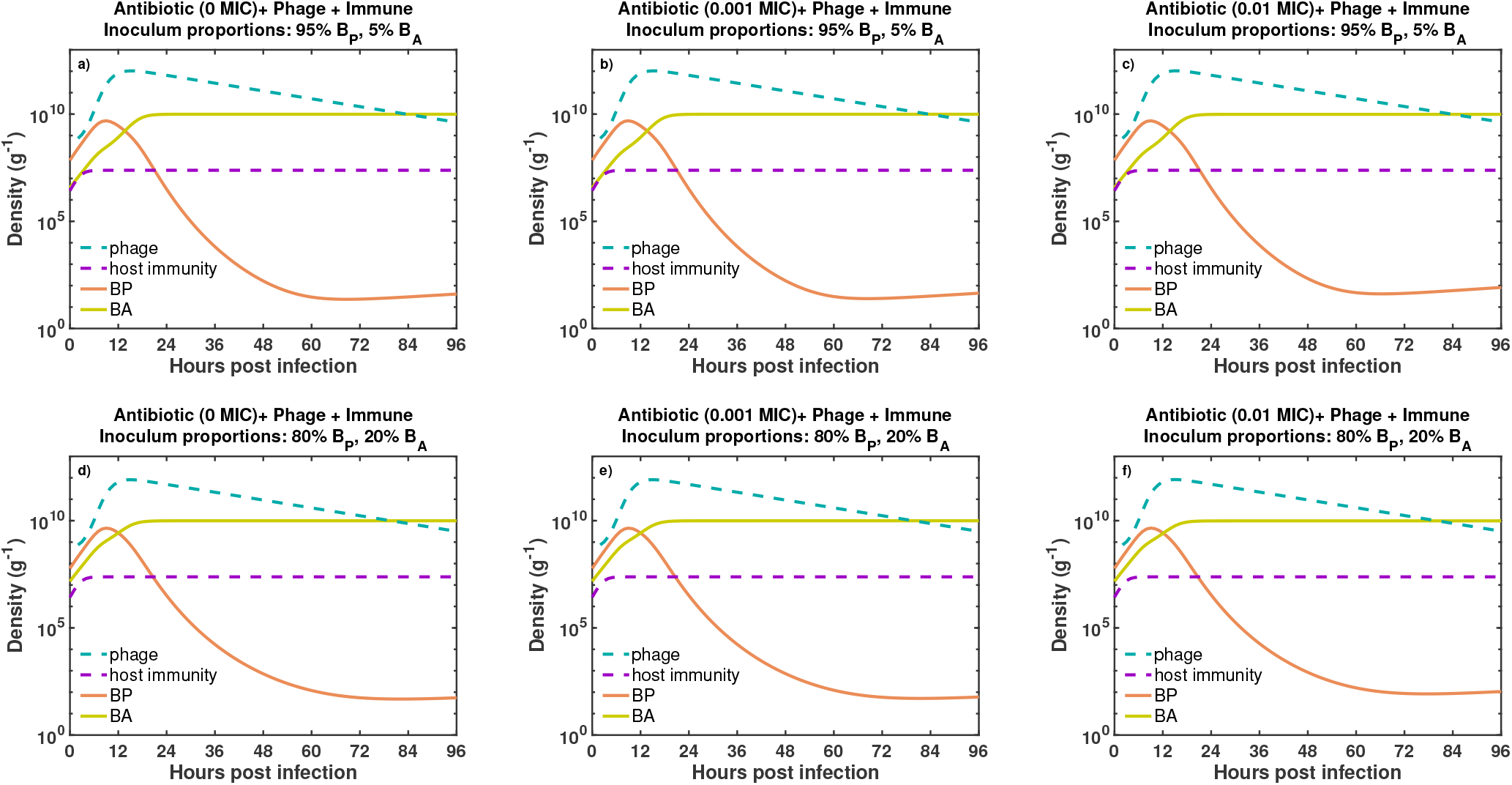
Bacterial dynamics given exposure to low levels of antibiotics. We simulate the effects of combination therapy plus innate immunity on inocula with non-trivial levels of *B*_*A*_. First, an inoculum composed of 95% *B*_*P*_and 5% *B*_*A*_ is treated with phage and different levels of antibiotic, 0, 0.001, and 0.01 X MIC (a, b, and c respectively). The same treatment is applied for an inoculum composed of 80% *B*_*P*_ and 20% *B*_*A*_ (d, e, and f, respectively). Initial bacterial and phage density are, *B*_0_ = 7.4 × 10^7^ CFU/g and *P*_0_ = 7.4 × 10^8^ PFU/g. Phage and antibiotic are administered two hours after infection. The simulation run was 96 hours (4 days).

**FIG. S2:**
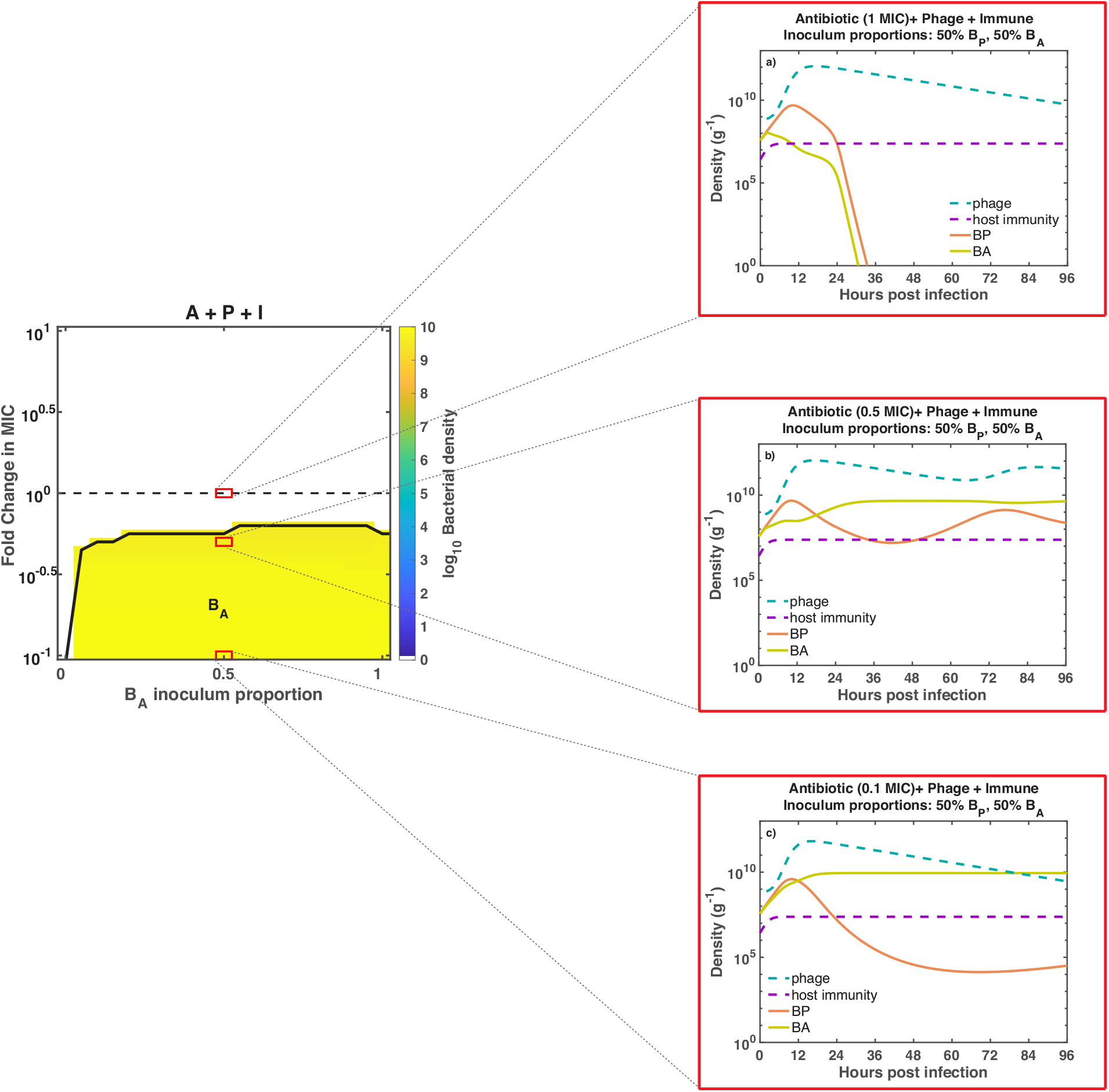
Accounting for variations in the concentration of antibiotic, from sub-MIC to MIC levels. We choose a particular inoculum composition (red boxes, 50%*B*_*P*_and 50%*B*_*A*_) from our heatmap and zoom in at the dynamics level. Bacterial dynamics correspond to different antibiotic levels: 1, 0.5, and 0.1 X MIC levels (a, b, and c, respectively). The colored areas on the heatmap indicate bacterial presence while the white areas indicate infection clearance after 96 hrs of treatment. Phage and antibiotic are administered two hours after infection. Initial bacterial and phage density, *B*_0_ = 7.4 × 10^7^ CFU/g and *P*_0_ = 7.4 × 10^8^ PFU/g, respectively.

**FIG. S3:**
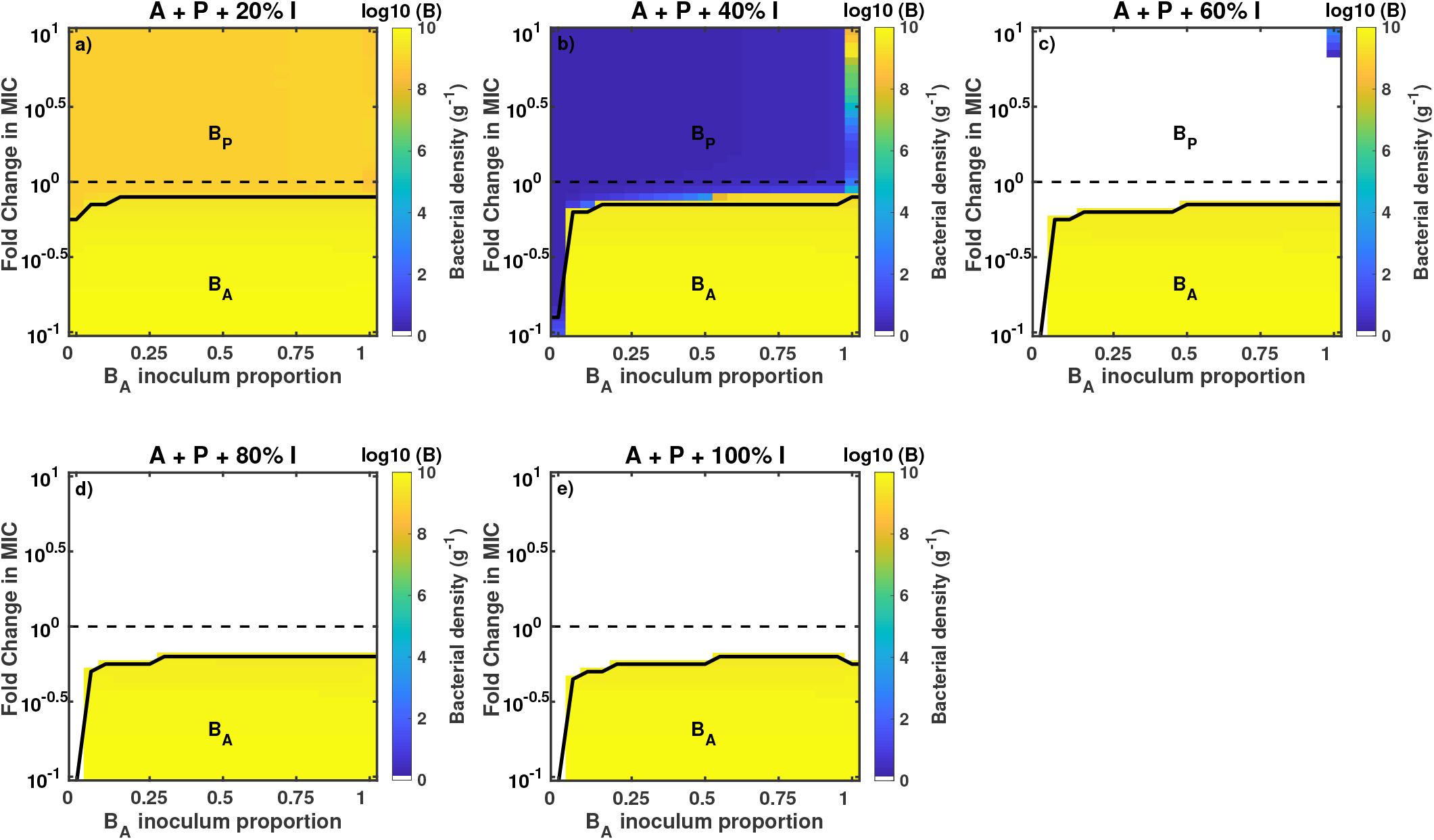
Bacterial density after 96 h of combined treatment with intermediate innate immunity levels. We extend our robustness analysis of Figure 5 (bottom) to account for intermediate levels of innate immune activation in the context of combined therapy. We vary the levels of innate immune response activation from 20% to 100% (a, b, c, d, and e). Bacterial density is calculated after 96 hr of treatment. Colored regions represent bacterial presence while white regions indicate infection clearance. Phage and antibiotic are administered two hours after infection. Antibiotic levels vary from 0.1 to 10 X MIC (MIC of ciprofloxacin = 0.014*µg/ml*). Initial bacterial and phage density are, *B*_0_ = 7.4 × 10^7^ CFU/g and *P*_0_ = 7.4 × 10^8^ PFU/g, respectively.

**FIG. S4:**
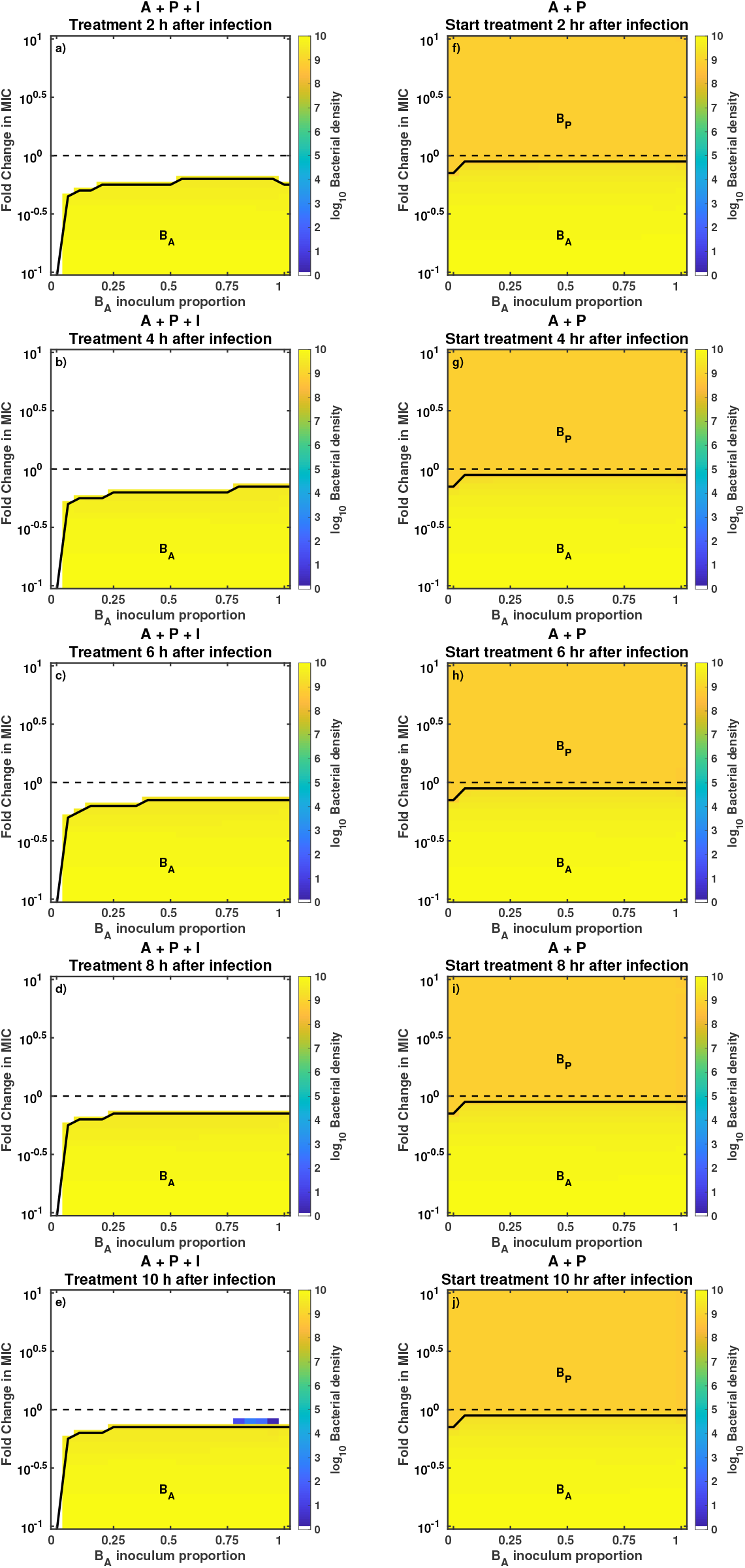
Time delays in the application of the combined treatment. We extend our robustness analysis of Figure 5 (bottom) to account for time delays in the start of the combined treatment. Phage and antibiotic were administered simultaneously 2, 4, 6, 8, and 10 hours after the beginning of the infection in the presence (Fig. S4 a-e) or absence (Fig. S4 f-j) of innate immunity. Colored regions on the heatmaps indicate bacterial presence while white regions indicate infection clearance. Antibiotic levels vary from 0.1 to 10 X MIC (MIC of ciprofloxacin = 0.014*µg/ml*). Initial bacterial and phage density are, *B*_0_ = 7.4 × 10^7^ CFU/g and *P*_0_ = 7.4 × 10^8^ PFU/g, respectively.

**FIG. S5:**
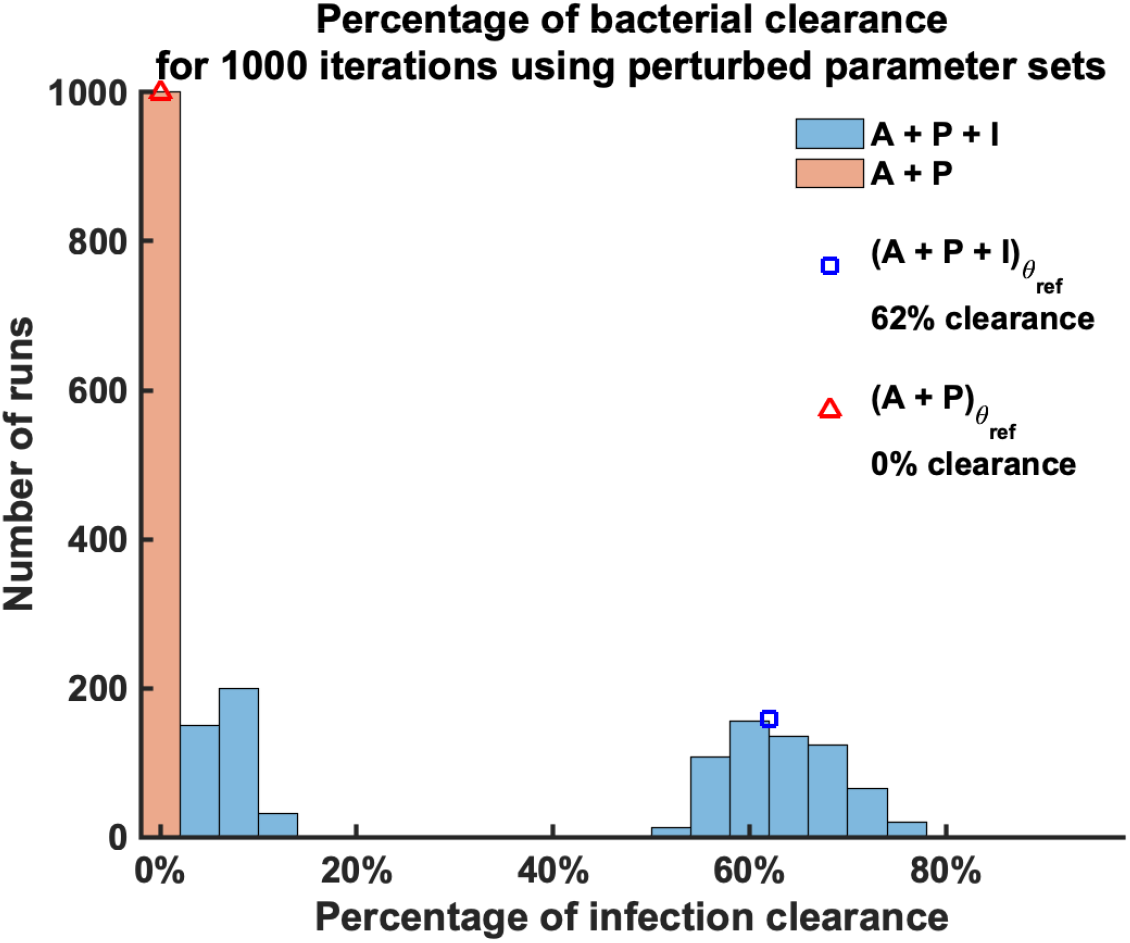
Parameter sensitivity analysis results. We show the distribution of the fraction of complete elimination for two therapeutic regimes A + P + I (blue) and A + P (red). We performed 1000 runs using perturbed parameter sets (*θ*_*per*_) and calculate the fraction of bacterial elimination for the two regimes. Moreover, we show the fraction of complete elimination for A + P + I (blue square) and A + P (red triangle) using the reference parameter set (*θ*_*ref*_).

**FIG. S6:**
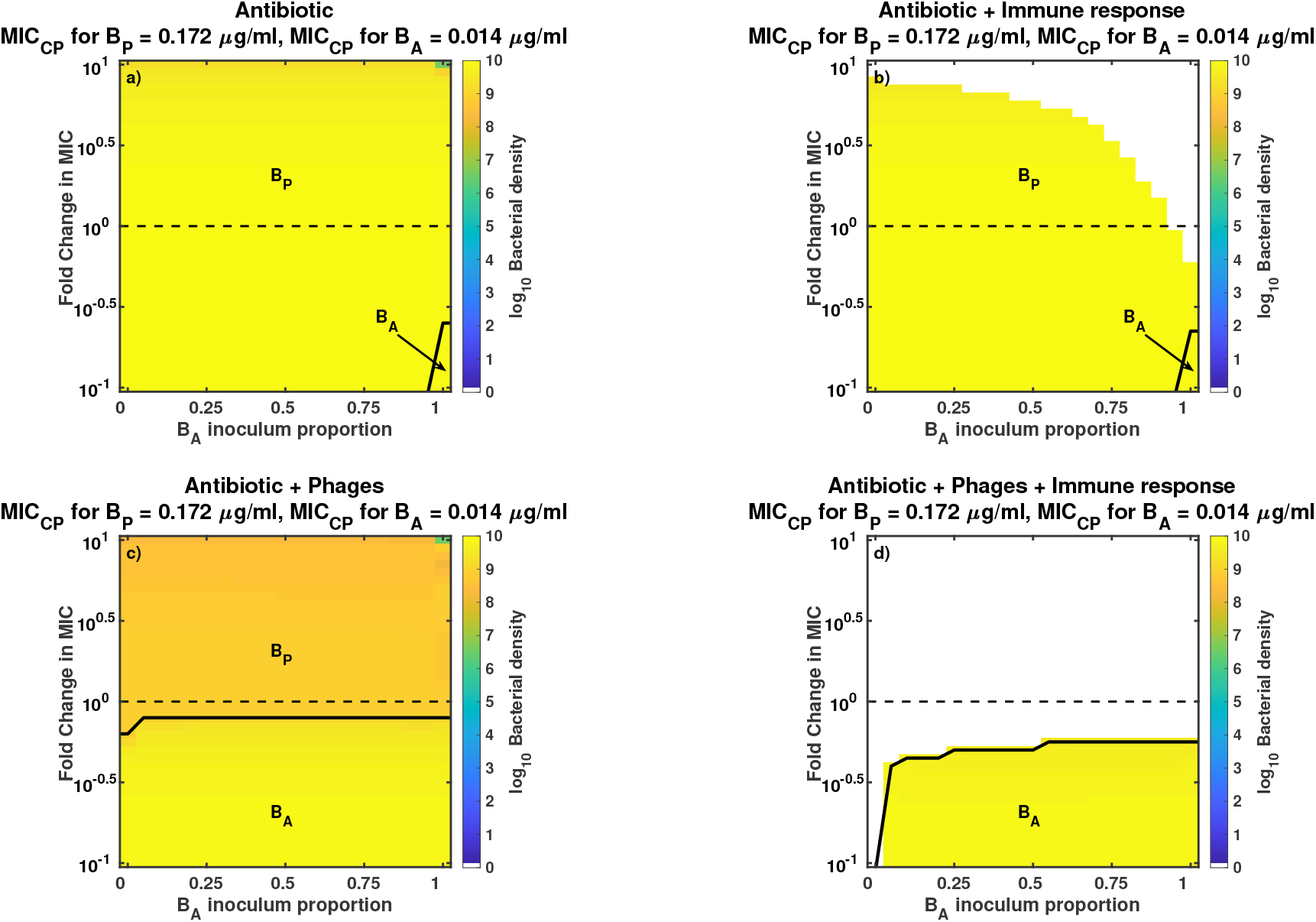
Robustness analysis of the partial resistance model for different antimicrobial strategies. Bacteria grew for 96 hours exposed to different antimicrobial strategies, antibiotic-only (a), antibiotic + innate immunity (b), phage and antibiotic (c), and phage-antibiotic combination plus active innate immunity (d). The heatmaps show the bacterial density at 96 hours post infection. Colored regions represent bacteria persistence while the clearance of the infection is represented by white regions. For the partial resistance model, phage can infect *B*_*A*_ with an infection constant of *ϕδ*_*P*_, being *δ*_*P*_ = 0.1. Moreover, the phage-sensitive strain, *B*_*P*_, is slightly sensitive to the antibiotic with an MIC of 0.172 *µ*g/ml. We simulate different concentrations of ciprofloxacin (CP); using the MIC of CP for the *B*_*A*_ strain as a reference (MIC = 0.014 *µ*g/ml). Fold changes in MIC go from 0.1 MIC (0.014 *µ*g/ml) to 10 MIC (0.14 *µ*g/ml). Furthermore, bacterial composition of the inoculum ranges from 100% phage-sensitive bacteria (0% *B*_*A*_) to 100% antibiotic-sensitive bacteria (100% *B*_*A*_). Initial bacterial density and phage density (c-d) are, *B*_0_ = 7.4 × 10^7^ CFU/g and *P*_0_ = 7.4 × 10^8^ PFU/g, respectively. Phage and antibiotic are administered two hours after the beginning of the infection.

**FIG. S7:**
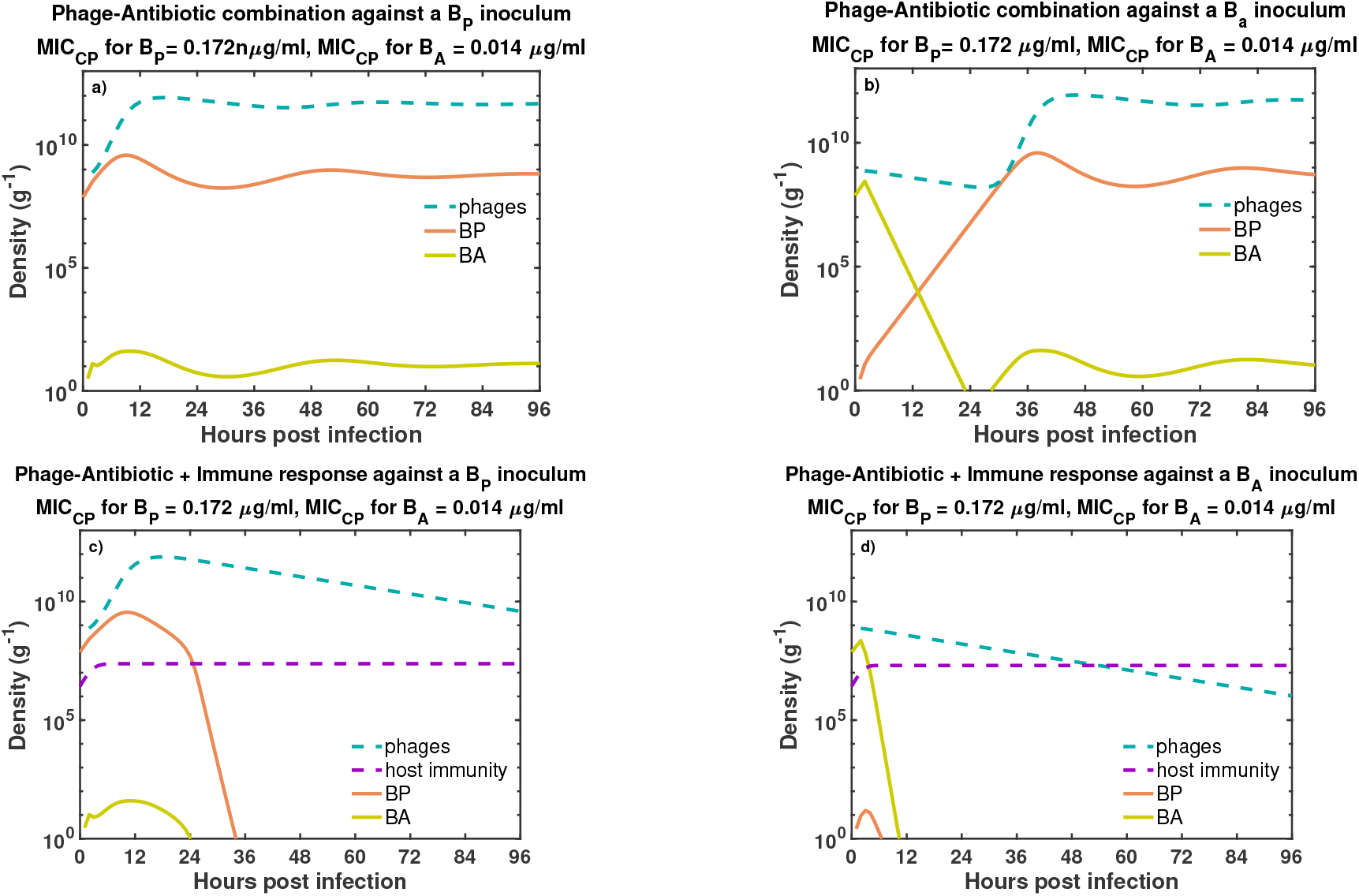
Time series of the partial resistance model in presence and absence of host immune response. We simulate the combination of phage and antibiotic against a phage-sensitive (a) and an antibiotic-sensitive (b) bacterial inoculum in the absence of the immune response. Moreover, we simulate a within-host scenario where the combined therapy interact with the immune response (purple dashed line) and phage-sensitive bacteria (c) or antibiotic-sensitive bacteria (d). Here, phage (blue dashed line) and antibiotic are administered two hours after the infection. Initial conditions are, *B*_0_ = 7.4 × 10^7^ CFU/g and *P*_0_ = 7.4 × 10^8^ PFU/g. The concentration of antibiotic (0.0350 *µg/ml* = 2.5×MIC for *B*_*A*_ strain) is maintained constant during the simulation, data not shown. The simulation run was 96 hours (4 days).

## Notes

#### Summary of Updates

In addition to minor changes to the text, we have made the following major changes: 1. Substantially expanded our discussion of the generality of the link between susceptibility to phage and antibiotic resistance mechanisms given binding to components of efflux systems, not only with P.a. but within other microbes as well. We hope that this discussion makes it self-evident that the modeling context is broadly relevant. 2. Included multiple new modeling analyses that confirm our core findings of an essential synergy between antibiotics, phage, and the immune system, while enhancing the robustness of the findings. To do so we evaluated the robustness of findings given variation in both the levels of antibiotics and state of the immune system as well as via a parameter sensitivity analysis.

